# A Programmable Optical Stimulator for the *Drosophila* Eye

**DOI:** 10.1101/147389

**Authors:** Xinping Chen, Walter D. Leon-Salas, Taylor Zigon, Donald Ready, Vikki Weake

**Affiliations:** Department of Biochemistry, Purdue University, 175 South University Street, West Lafayette, Indiana, USA; School of Engineering Technology, Purdue University, 401 North Grant Street, West Lafayette, Indiana, USA; Department of Biological Sciences, Purdue University, West Lafayette, Indiana, USA; Purdue Center for Cancer Research, 201 South University Street, West Lafayette, Indiana, USA

**Keywords:** optical stimulation, light-emitting diodes, embedded computers, Drosophila, Rhodopsin, open hardware

## Abstract

A programmable optical stimulator for *Drosophila* eyes is presented. The target application of the stimulator is to induce retinal degeneration in fly photoreceptor cells by exposing them to light in a controlled manner. The goal of this work is to obtain a reproducible system for studying age-related changes in susceptibility to environmental ocular stress. The stimulator uses light emitting diodes and an embedded computer to control illuminance, color (blue or red) and duration in two independent chambers. Further, the stimulator is equipped with per-chamber light and temperature sensors and a fan to monitor light intensity and to control temperature. An ON/OFF temperature control implemented on the embedded computer keeps the temperature from reaching levels that will induce the heat shock stress response in the flies. A custom enclosure was fabricated to house the electronic components of the stimulator. The enclosure provides a light-impermeable environment that allows air flow and lets users easily load and unload fly vials. Characterization results show that the fabricated stimulator can produce light at illuminances ranging from 0 to 16000 lux and power density levels from 0 to 7.2 mW/cm^2^ for blue light. For red light the maximum illuminance is 8000 lux which corresponds to a power density of 3.54 mW/cm^2^. The fans and the ON/OFF temperature control are able to keep the temperature inside the chambers below 28.17°C. Experiments with white-eye male flies were performed to assess the ability of the fabricated simulator to induce blue light-dependent retinal degeneration. Retinal degeneration is observed in flies exposed to 8 hours of blue light at 7949 lux. Flies in a control experiment with no light exposure show no retinal degeneration. Flies exposed to red light for the similar duration and light intensity (8 hours and 7994 lux) do not show retinal degeneration either. Hence, the fabricated stimulator can be used to create environmental ocular stress using blue light.

Specifications table

Specifications table
|  |  |
| --- | --- |
| Hardware name | Drosophila Optical Stimulator |
| Subject area | Biological Sciences |
| Hardware type | Electrical engineering and computer science |
| Open Source License | CERN Open Hardware License v1.2 |
| Cost of Hardware | USD 757.04 |
| Source File Repository | Google Drive |

## 1. Hardware in Context

Light is essential for vision, and is sensed by a specialized light-sensitive layer of tissue in the eye, the retina, that sends visual information through the optic nerve to the brain. Notably, light functions as both a signal and as an environmental stress that can induce retinal degeneration under specific conditions or in particular genotypes [1]. Specialized neurons, called photoreceptor neurons, sense light in the retina and convey information to the brain. Given that photoreceptor neurons have the same lifespan as the organism and do not regenerate, cumulative exposure to light occurs with age. Light exposure is required for vision, but because it can induce retinal degeneration, it also plays a role in eye disease, a role that is more pronounced as the eye ages.

Advanced age is a major risk factor for ocular diseases such as age-related macular degeneration, glaucoma, and diabetic retinopathy [2, 3, 4]. The incidence of blindness and low vision increases rapidly with age, and persons older than 80 years have incidences of blindness that range from 1.8 - 6.8% depending on ethnicity [5]. Defined model systems are required to study how factors including genotype, environmental stress and age interact to increase the susceptibility of the eye to ocular disease. The fruitfly *Drosophila melanogaster* provides a powerful model system in which to study how environmental stress contributes to age-related visual decline. The appeal of the fruitfly to these studies comes from its short life-span, amenability to genetic manipulation, and susceptibility to specific wavelengths of light, which induce photoreceptor degeneration in a controlled manner.

One standard technique employed to induce retinal degeneration in fruitflies involves exposing flies to high-intensity blue light for extended periods of time. In [6] Cosens showed that prolonged exposure to blue wavelengths of light, but not other wavelengths, severely disrupts the rhabdomere structure of retinula (R) cells in white-eyed flies. In these pioneering studies, the authors exposed 6 to 12 individual white-eyed flies to blue light with an intensity of 0.1 mW/cm^2^ for 22 hours in glass tubes and to blue light with intensity 2.0 mW/cm^2^ for 0.5 to 12 hours in Petri dishes. Blue light was generated using tungsten bulbs and optical band-pass filters. Subsequent studies determined that the threshold for blue light-induced retinal degeneration in white-eyed flies was 20 log quanta/cm^2^ for single flies subjected to intense light stimulus from a 100 W Mercury vapor arc lamp for 30 s [7]. In [8] pulsed blue light was used in retinal stimulation studies. Blue light was generated using a 300 W xenon/mercury lamp and a optical band-pass filter.

The development of blue light-emitting diodes (LEDs) provided a new light source with high efficiency, narrow emission spectra and small form factor. Blue LEDs have been used to optically stimulate retinal cells of flies [9] and mice [10, 11, 12]. In each of these studies, custom-made LED-based stimulators were fabricated and used. However, these stimulators had only basic functionality (lights turning on and off), lacked programmability and monitoring and only a small number of flies could be subjected to light stress each time. Moreover, these stimulators only emitted light of a single color (blue). It has been shown that orange and red light can have the opposite effect of blue light, that is, regeneration of light-sensitive proteins, such as Rhodopsin, in the six outer photoreceptor neurons (R1-R6). Hence, red light can be used in control experiments to verify that light-induced retinal degeneration is indeed wavelength dependent and to regenerate light-sensitive proteins. Hence, a desirable feature in an optical stimulator would be to emit light in blue and red colors. The hardware design files and source code of previously published optical stimulation devices have not been made available to the public. Moreover, due to the small market size for such devices, commercial solutions are not available. Hence, new researchers needing to carry out similar experiments are faced with the challenge and time-consuming task of developing their own devices from scratch.

This paper presents the design, fabrication and test of a programmable dual-color LED-based device suitable for optical stimulation of fruitflies. Design files for the hardware and software components are openly shared so that an identical device can be built by researchers needing to perform controlled optical stimulation experiments. Our optical stimulator was used in several experiments aimed at inducing retinal degeneration in fly photoreceptor cells in a controlled manner to obtain a reproducible system for studying ocular stress. The device can generate blue light at intensities ranging from 0 to 16,000 lux and optical power densities up to 7.2 mW/cm^2^. The device parameters such as stimulation time, light intensity and color (red or blue) can be programmed locally or remotely through a menu-driven user interface. Stimulation can be performed simultaneously on two independent chambers. Both chambers are equipped with light and temperature sensors to monitor the stimulation conditions, which are automatically recorded in computer files. The chambers are also equipped with fans and an ON/OFF temperature controller to ensure that temperature inside the chambers stays below 28°C. The device has been characterized and used to establish a protocol for assessing blue light-induced retinal degeneration in young fruitflies. Experimental results show retinal degeneration in flies exposed to blue light at ~7900 lux for 8 hours. However, flies exposed to red light of similar intensity and duration showed no retinal damage. Hence, the presented stimulator can be used to create environmental ocular stress using blue light.

### 1.1. Background

*Drosophila* have eyes that are composed of approximately 750 units known as ommatidia (Fig. 1). Each ommatidium contains 20 cells including 8 photoreceptor neurons called R cells (R1 – R8). Each R cell expresses a membrane-localized protein, Rhodopsin, which functions to sense light. Rhodopsins are highly conserved as light-sensors in both insects and in vertebrates. Rhodopsin 1 (Rh1) is the major Rhodopsin in Drosophila and is expressed in a subset of photoreceptors neurons; R1 – R6 [13, 14]. Rh1 is localized predominantly in rhabdomeres, which are specialized light-sensing organelles on the apical surface of each photoreceptor neuron. Absorption of a blue light photon (460 – 490 nm) by Rh1 causes a conformational change to the active form of the receptor protein, metarhodopsin (metaRh1), which signals through a G-protein-coupled cascade to trigger the opening of the transient receptor potential (TRP) channel, influx of calcium, and depolarization of the photoreceptor cell [15]. An orange or red photon is required to convert metaRh1 back to Rh1 [16]. Thus, prolonged exposure to blue light causes endocytosis and degradation of metaRh1 since the metaRh1 cannot be converted back to Rh1 [17, 9]. In contrast, under normal white light conditions, metaRh1 is readily converted back to Rh1. Both the calcium influx and endocytosis of metaRh1 caused by prolonged exposure to blue light induce retinal degeneration in *Drosophila* photoreceptors [8, 18]. Thus, exposing *Drosophila* to blue light provides a controlled system to study the process of light stress-induced retinal degeneration and to identify factors that protect photoreceptors from damage.

**Figure 1:**
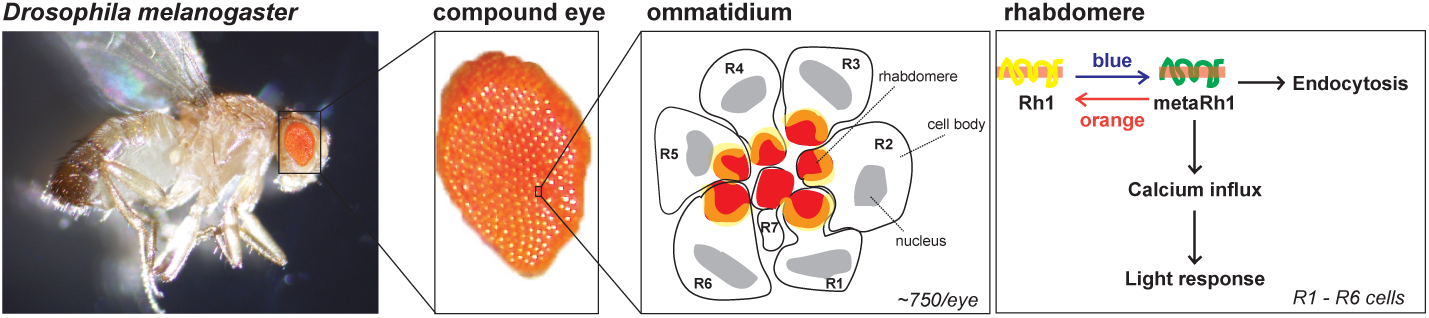
Structure of a Drosophila compound eye. The fruitfly *Drosophila melanogaster* has a compound eye composed of ~750 units known as ommatidia. Each ommatidium contains cone cells, pigment cells, bristle cells and 8 different photoreceptor neurons; R cells 1 – 8. The R1 – R6 photoreceptors express the light-receptor Rhodopsin 1 (Rh1; shown in yellow) and occupy the outer region of the ommatidium, while R7 and R8 photoreceptors are present in the center of the ommatidium. Since R7 and R8 photoreceptors are stacked on top of one another, only R7 (shown here) or R8 can be observed in a given cross-section image of each ommatidium. Photoreceptor neurons in flies have a characteristic light-sensing organelle, the rhabdomere (shown in red), that is composed of an ordered array of tiny membrane projections called microvilli. Light is sensed in rhabdomeres of R1 – R6 cells by Rh1, which absorbs a blue light photon (460 – 490 nm) inducing a conformational change to metarhodopsin (metaRh1). This conformational change can be reversed by absorption of an orange or red photon, and this process usually occurs rapidly in the presence of white light. However, prolonged exposure to blue light in the absence of orange or red wavelengths results in accumulation of metaRh1, inducing calcium influx into the photoreceptor and continual light signaling. Over time, this continual light signaling in the presence of blue light can be downregulated by removing metaRh1 through the process of endocytosis, in which the plasma membrane containing the receptor is engulfed into the interior of the cell, thus removing metaRh1 from the rhabdomere. Photoreceptors R7 and R8 express different Rhodopsin proteins and respond to different wavelengths of light.

## 2. Hardware Description

This section describes the design and fabrication process of the programmable optical stimulator for *Drosophila*. The design process started with a list of technical requirements set by the target application and feedback from end users regarding user interface and functionality. Based on these requirements an architecture for the stimulator was chosen. Specific components and devices were then sized and selected.

### 2.1. Design Requirements

The following design requirements were identified:

1. To increase the number of flies that can be exposed to light stress in each experiment, the stimulator should be able to work with standard fly vials that can hold between 50 and 100 flies. The standard transparent plastic vials used in *Drosophila* experiments have a height of 95 mm high and a diameter of 25 mm.
2. The stimulator should have light emission in two different wavelengths: 460 nm to 470 nm (for blue color light) and 620 nm to 630 nm (for red color light) [19].
3. The stimulator should emit light with an intensity and duration such that the threshold for light-induced retinal degeneration of 20 log quanta/cm^2^ [7] is met or exceeded.
4. The user should be able to vary the light intensity from maximum intensity to zero intensity.
5. The device should be able to stimulate two vials simultaneously. The stimulation parameters for each vial, such as color, duration and intensity, should be set independently for each vial. The second vial will be used to run parallel or control experiments. Control experiments with opposite or no stimuli are necessary to isolate the effects that wavelength, light intensity and stimulation duration might have on the phenotype being assessed.
6. The stimulator should have a light sensor to provide feedback to the user and to monitor light stability during prolonged experiments.
7. Temperature must be monitored and controlled within 22°C and 28°C to prevent heat-induced stress response [20, 21].
8. The stimulator should have an easy-to-use interface that allows users to set stimulation parameters and to start and stop a stimulation session. It is highly desirable to have the ability to remotely initiate, monitor and terminate stimulation sessions. A stimulation session typically runs for 8 to 12 hours. Hence, it is expected that users would like to monitor an experiment remotely from home or from a place outside the lab. In this regard, it will be advantageous if the stimulator can connect to the network infrastructure already present in a typical lab such as Ethernet and WiFi networks.
9. It is desirable that stimulation parameters as well as the periodic readings from the light and temperature sensors are recorded electronically. This recorded information will be used for archival purposes, calibration and to determine if light and temperature levels are kept within expected ranges within a stimulation session.
10. Each chamber should be impermeable to external light source s.

### 2.2. Design Process

#### 2.2.1. Embedded Computing Platform

The required degree of configurability, user interface and data loging suggests the use of an embedded computing solution. The hardware architecture of the embedded computing solution is shown in Fig. 2. It consists of an embedded single-board computer, two independent stimulation chambers (left and right), an LCD screen, keyboard, mouse and a power supply. The LCD screen, keyboard and mouse provide a familiar and an easy-to-use interface. Each chamber contains several blue and red LEDs, LED driver circuits and a light and a temperature sensors. Each chamber can stimulate one vial at a time.

**Figure 2:**
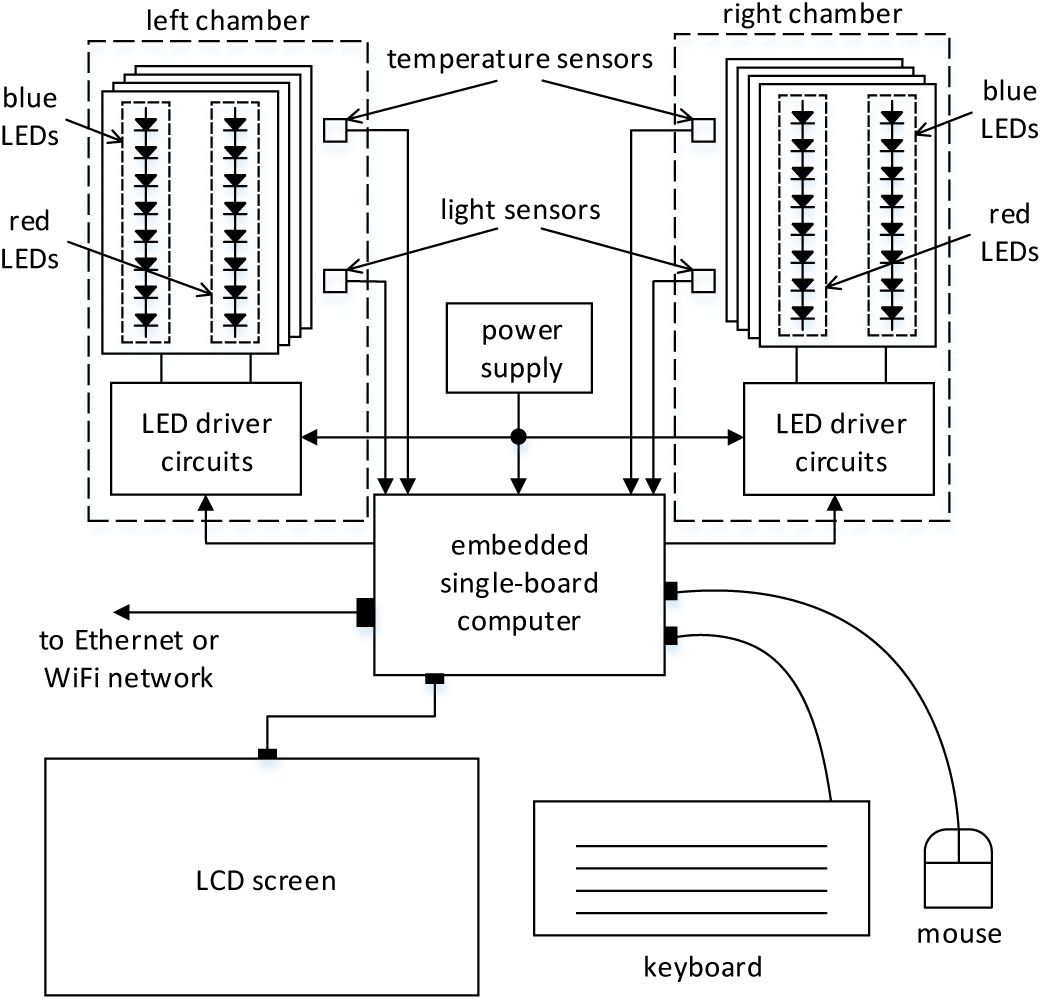
Hardware architecture of the programmable optical stimulator. The hardware of the stimulator comprises a single-board Linux embedded computer, a power supply, an LCD screen, a keyboard and a mouse and two stimulation chambers. Each chamber contains several red and blue LEDs with their corresponding driver circuits.

Using an embedded computer with an Operation System (OS), such as Linux, with drivers for many input/output peripherals, simplifies the implementation of connectivity over Internet Protocol (IP) networks, data logging on electronic files and the creation of a user interface. Linux, being a popular open-source OS, enjoys a wide support community that the designer can tap into to accelerate development time. Several single-board computers with Linux support are commercially available [22]. The Raspberry Pi has the right balance of CPU speed, peripherals, low cost and size. Hence, it was chosen as the embedded computing platform for the optical stimulator.

#### 2.2.2. LED Array

In order to generate light with sufficient illuminance or power density to trigger retinal degeneration, each stimulation chamber is equipped with several blue LEDs. The 5050-B1200 blue LEDs from Superbrightleds [23] were chosen as the light sources in the programmable optical stimulator. The 5050-B1200 LEDs are low-cost and widely available LEDs and come in a convenient surface mount package of size 5 mm × 5 mm × 1.6 mm. The 5050-B1200 is a multi-die LED, that is, it comprises three independent LED dies in a single package that collectively provide a nominal luminous intensity of 1.2 candela (cd) at a forward current of 20 mA through each die. The nominal wavelength of the emitted light for this LED is 465 nm and its 50% viewing angle is 120°.

The illuminance or luminous flux per unit area of these LEDs, *E_v_*, can be calculated as follows [24]:

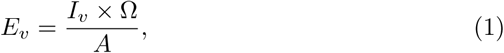

where, *A* is the illuminated area and *I_v_* is the luminous intensity across the solid angle Ω. *E_v_* is expressed in lux when *I_v_* is expressed in candela (cd) and Ω in steradians (sr) and *A* in square meters. The solid angle can be estimated from the 50% viewing angle (*θ*) as follows:

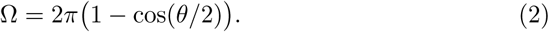

Replacing (2) in (1) yields the following expression for the illuminance:

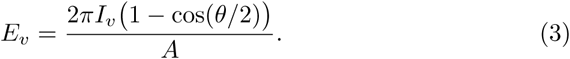

The power density or power per unit area emitted by a single LED, *P_led_*, can then be calculated from the illuminance using the following equation:

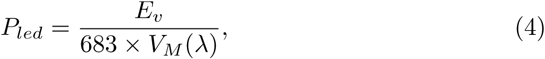

where, *V_M_* (*λ*) is the spectral luminous efficiency function for photopic vision. This function describes the average spectral sensitivity of the human visual system. It has a maximum of 1.0 at a wavelength of 555 nm. At a wavelength of 465 nm, *V_M_* (*λ*) = 0.0818. Replacing (3) in (4) yields:

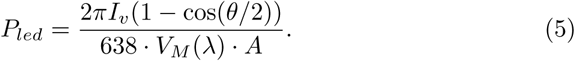

For the selected LEDs, *I_v_* = 1.2 cd and *θ* = 120°. Considering a cylindrical chamber with 9.5 cm of internal diameter and a height of 12 cm, the illuminated area becomes 358.14 cm^2^. Hence, *P_led_* = 188.3 *μ*W/cm^2^. The luminous flux is defined as *Φ_v_* = *I_v_* × Ω. Hence, the luminous flux of a single LED can be estimated to be 3.77 lumen (lm). The number of blue LEDs needed in each chamber can be estimated from the threshold for retinal degeneration.

Previous studies have established that the threshold for retinal degeneration is 20 log quanta/cm^2^ [7]. A log quanta/cm^2^ is defined as follows:

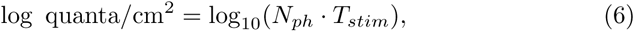

where, *N_ph_* is the number of photons per unit area per unit time and *T_stim_* is the stimulation time. *N_ph_* can be estimated using:

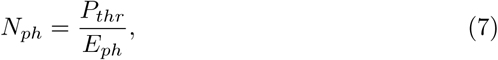

where, *P_thr_* is the minimum power per unit area that causes retinal degeneration in *T_stim_* time. *E_ph_* is the energy per photon and is equal to *hc/*λ, where *h* is Planck’s constant, *c* is the speed of light and λ is the photons’ wavelength. Replacing (7) in (6) and solving for *P_thr_* yields:

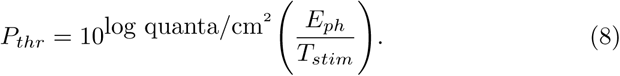

The number of blue LEDs, *K*, needed in each chamber to achieve the threshold for retinal degeneration is therefore equal to:

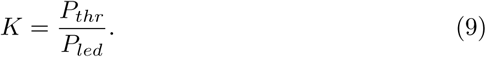

Considering one hour to be the shortest stimulation time, photons with a wavelength of 465 nm and 20 log quanta/cm^2^, equation (8) gives *P_thr_* = 11.9 mW/cm^2^. Hence, the required number of blue LEDs is 63.05. In the fabricated programmable optical stimulator, a total of 64 of these LEDs per chamber are employed. An equal number of red multi-die LEDs (5050-R1200) are employed. The 5050-R1200 comprises three red LED dies in a single 5 mm × 5 mm × 1.6 mm package. This LED has a nominal luminous intensity of 1.2 cd at 20 mA, an emission centered at 625 nm and a 50% viewing angle of 120° [25]. Hence, each chamber contains 128 blue and red LEDs.

#### 2.2.3. Circuit Design

The LEDs are grouped in groups of 16 LEDs (8 blue and 8 red) and assembled on custom printed circuit boards (PCBs) called LED boards. Each LED board also includes an LED driver circuit and a power and data bus. To uniformly illuminate a fruitfly vial, eight LED boards are arranged concentrically inside a stimulation chamber around the vial. Hence, a vial is illuminated by 64 multi-die blue LED dies and 64 multi-die red LED dies. This setup, shown in Fig. 3, is repeated for the left and right chambers. Figure 3 also shows the schematic diagram of the circuitry in each LED board along with other supporting circuits such as a voltage regulator and a general purpose input/output (GPIO) expander.

**Figure 3:**
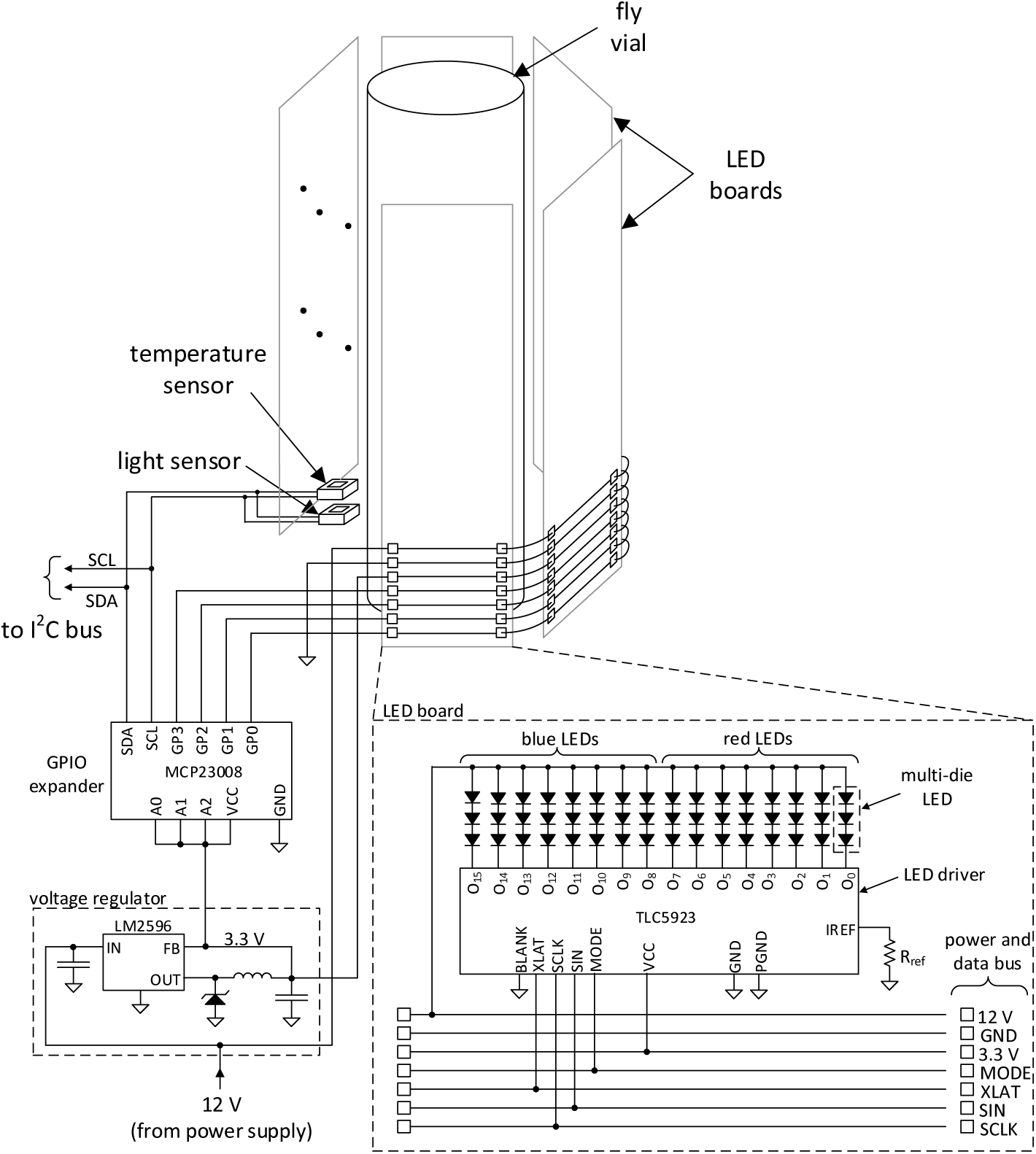
Schematic diagram of the electronic circuit employed to control the current through the LEDs and to monitor light and temperature inside a chamber. A 16-channel LED driver with serial interface enables independent LED current control in 128 discrete steps. Each LED board contains eight multi-die blue LEDs and eight multi-die red LEDs. Eight LED boards surround the fly vial to provide uniform illumination. The LED boards have a modular design that allows them to share a common power and data bus and minimize wiring. The temperature and light sensors, used to monitor the condition inside the chamber, connect directly to the I^2^C bus of the embedded computer.

The LED board is designed such that the individual LED dies in each multi-die LED are connected in series. The LED driver is the integrated circuit TLC5923 from Texas Instruments. This circuit can independently control the current of 16 LEDs from 0 to a maximum of *I_max_*. The current *I_max_* is set by resistor *R_ref_* as follows [26]:

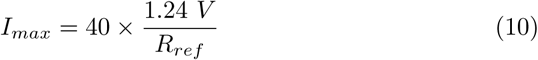

Using (10), an *I_max_* of 20 mA is obtained with a resistor of 2.48 kΩ. In practice, the closest resistor value commercially available in a surface mount package is 2.49 kΩ with 1% tolerance. Employing this resistor results in a maximum current of 19.9 mA through each LED die. The TLC5923 has a digital serial interface, comprised of signals MODE, XLAT, SIN and SCLK, that allows a computer to set the current in each one of its outputs (O_0_ to O_15_) to a value from 0 to *I_max_* in 128 steps. This feature is used in our design to allow the user to vary the LEDs’ current, and consequently their luminous intensity, from 0% (totally off) to 100% (fully on with maximum light emission).

The blue LEDs dies have a forward voltage drop of 3.25 V for a current of 20 mA. Given that the three blue LED dies in each multi-die LED are connected in series, a power supply voltage of 9.75 V or higher is required. In the programmable optical stimulator, a 12 V power supply is employed since high-current 12 V supplies are widely available.

The LED boards have a modular design that allows the embedded computer to control several boards with minimum wiring by sharing a common power and data bus that carries power signals (12 V and 3.3 V) and serial control signals (MODE, XLAT, SIN and SCLK). The MCP23008 GPIO expander is employed to further simplify wiring to and from the embedded computer. The port expander connects to the Inter-Integrated Circuit (I^2^C) serial bus of the embedded computer which is also shared by the light and temperature sensors. The temperature sensor employed is the MCP9808 from Microchip. This sensor can measure temperature in the −40°C to +125°C with an accuracy of 0.25°C [27], which is sufficient for the requirements of our application. The light sensor is the TSL2561 from TAOS. This sensor has a broadband photodiode sensitive to visible and infrared light and a second photodiode sensitive only to infrared light. The current output of these two photodiodes are converted to digital and sent to the embedded computer over the I^2^C bus where an empirical formula, that approximates the spectral luminous efficiency function for photopic vision is used to calculate the illuminance in units of lux [28].

The voltage regulator steps down the 12 V from the power supply to the 3.3 V required by the LED driver. Given that either the blue or the red LEDs are turned on at any given time, the maximum current drawn by the LEDs in one chamber is 20 mA/LED×8 LEDs/board×8 boards/chamber=1.28 A. Thus, the two chambers draw a maximum current of 2.56 A. Considering an additional 3 A to power the embedded computer, LCD screen and peripherals, yields a total current consumption of 5.56 A. An off-the-shelf 12 V, 72 W power provides enough current to power up the programmable optical stimulator.

Not all the power delivered by the power supply gets converted to light. A portion of this power is converted to heat by the LEDs and the LED driver circuits. The amount of heat produced by these elements can be estimated from the power delivered by supply, the luminous flux generated by the LEDs and the luminous efficacy of the LEDs. The maximum power delivered by the supply to the LEDs to one chamber is 12 V× 1.28 A = 15.4 W. The luminous flux of a single multi-die LED was calculated to be 3.77 lm. Hence, the luminous flux of 64 multi-die LEDs in one chamber is 241.28 lm. Assuming a luminous efficacy of 60 lm/W of a typical LED, the amount of power that is actually converted to light is 4.02 W. Hence, from the 15.4 W delivered by the supply to each chamber, 11.38 W get converted to heat. It should be noted that is the maximum amount of heat generated and corresponds to the LEDs emitting maximum luminous intensity. Over time this much heat will raise the temperature inside the chambers to levels that will induce the heat shock stress response in flies. This is undesirable for the experiments since it could interfere with the analysis of retinal degeneration and/or cause the flies to die.

To keep the temperature inside the chambers within an acceptable range, active cooling using forced air flow is used. The required air flow, *F_a_*, to keep the temperature inside the chamber below *T_o_*_°_C can be calculated using the following equation [29]:

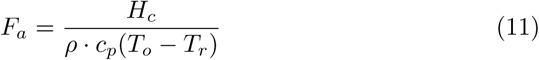

where, *H_c_* is the heat generated inside the chamber in Watts, *ρ* is the density of air, *c_p_* is the specific heat of air and *T_r_* is the air temperature outside the chamber (typically room temperature). Replacing the numerical values of *H_c_* = 11.38 W, *ρ* = 1.2 kg/m^3^, *c_p_* = 1005 J/kg K, *T_o_* = 28°C and *T_r_* = 23° C in (11), yields an air flow rate of 1.57 × 10^−3^ m^3^/s or 4 cubic feet per minute (CFM). An off-the-shelf fan with an air flow of 5.7 CFM is used and provides enough air flow to cool down the air in each chamber. When the LEDs are not working at maximum intensity, less heat is generated and the fans should be turned off to prevent the temperature inside the chambers to drop below about 20 °C. Extreme temperatures (both high and low) can induce stress responses in *Drosophila* [30]. To accomplish this requirement, each fan is independently controlled by the embedded computer via a MOSFET transistor. When the temperature inside a chamber exceeds 24.5°C, the embedded computer turns on the corresponding fan. Likewise, if the temperature drops below 23.5°C, the fan is turned off.

#### 2.2.4. Enclosure

An custom enclosure to house the different components of the programmable optical stimulator was fabricated. The enclosure was fabricated using 3D printed parts and laser-cut acrylic pieces. A lid on each chamber permits loading and unloading of fly vials while blocking ambient light during stimulation. The chamber design allows air to flow underneath the lid through holes under the lid. Air flow can bend and pass through the holes but light can not. This detail is depicted in Fig. 4 which shows the enclosure through a cut-through plane so that the chamber design details can be observed. The blue arrows indicate air flow.

**Figure 4:**
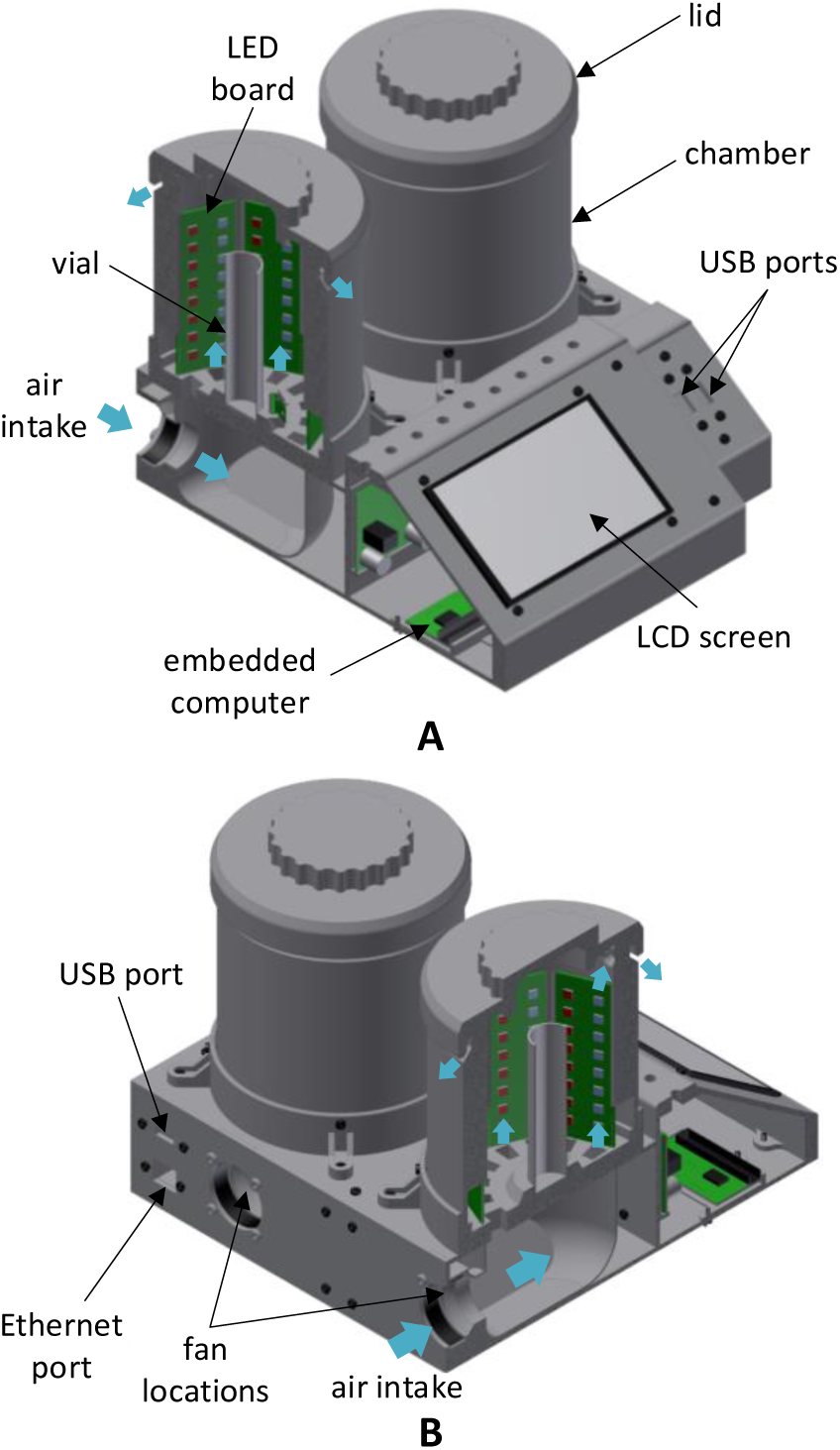
Fabricated enclosure. The enclosure was fabricated using 3D printed and laser-cut acrylic parts. The design allows air flow in and out of the chamber while blocking ambient light. The blue arrows indicate air flow. (A) Front view with cut-through plane. (B) Rear view with cut-through plane.

A five-inch LCD display with 800×480 pixel resolution serves as the screen of the embedded computer. Two USB connectors on the front and one on the back of the enclosure provide connectivity for a keyboard, mouse and a WiFi dongle. An Ethernet connector is installed on the back of the enclosure to allow connection to an Ethernet network.

#### 2.2.5. Software

A computer program that controls the LED brightness, reads the sensors’ outputs, interacts with the user and records events was developed. This program was written in the Python programming language and runs on the embedded computer. The user interface was created using `curses`, which is a programming library that allows the development of text-based user interfaces. A text-based user interface has the advantage of being simple to use and able to run from a Windows console or a Telnet or SSH client which makes it easier to operate remotely. The user interface is menu driven and allows the user to set stimulation parameters, start and stop a stimulation session and monitor illuminance and temperature independently for each chamber. The stimulation parameters that can be set by the user are: duration, light color and light intensity in a relative scale from 0% to 100% where 100% represents maximum current (*I_max_*) flowing through the LEDs. Figure 5 shows a screen capture of the user interface.

**Figure 5:**
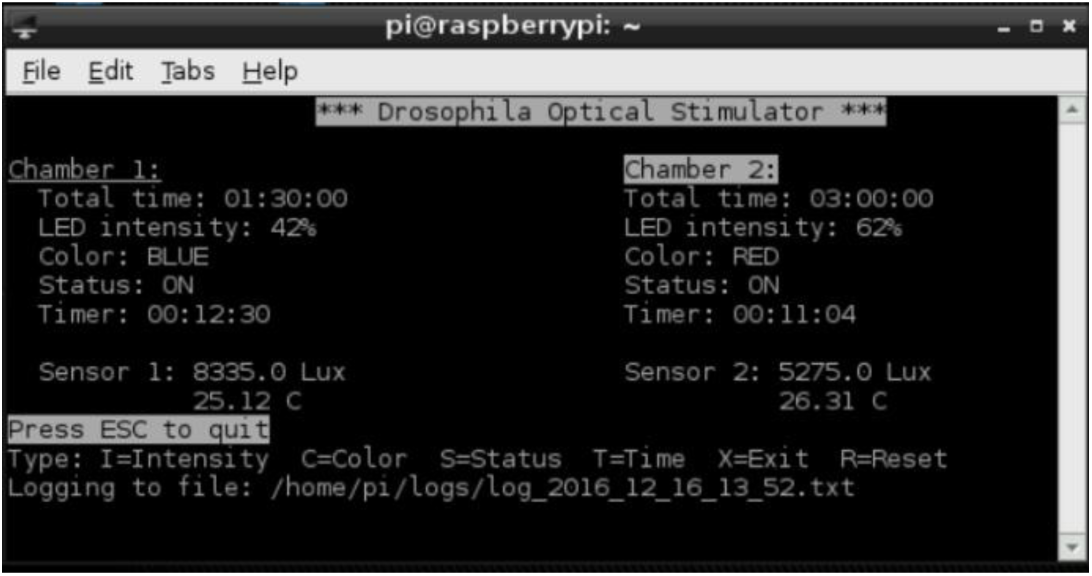
Screen capture of user interface. The user interface is a text-based interface that allows the user to start and stop a stimulation session as well as change stimulation parameter such as light color, light intensity and stimulation time.

A flow diagram of the program is shown in Fig. 6. The program consists of a main loop that reads and displays the outputs of the light and temperature sensors for each chamber. Every few seconds the sensors readings are recorded in a text log file for archival and troubleshooting purposes. Other events such as when LEDs are turned on or off, or when the light intensity is changed by the user, or expiration of the stimulation time are also recorded in the log file. Once a stimulation session has started, the program keeps track of the LED on time and compares it with the stimulation time selected by the user. When the on time exceeds the selected stimulation time, the LEDs are turned off. The program also implements an ON/OFF temperature control as follows: when the temperature inside the chamber exceeds 24.5°C, its fan is turned on, when the temperature inside the chamber drops below 23.5°C, its fan is turned off.

**Figure 6:**
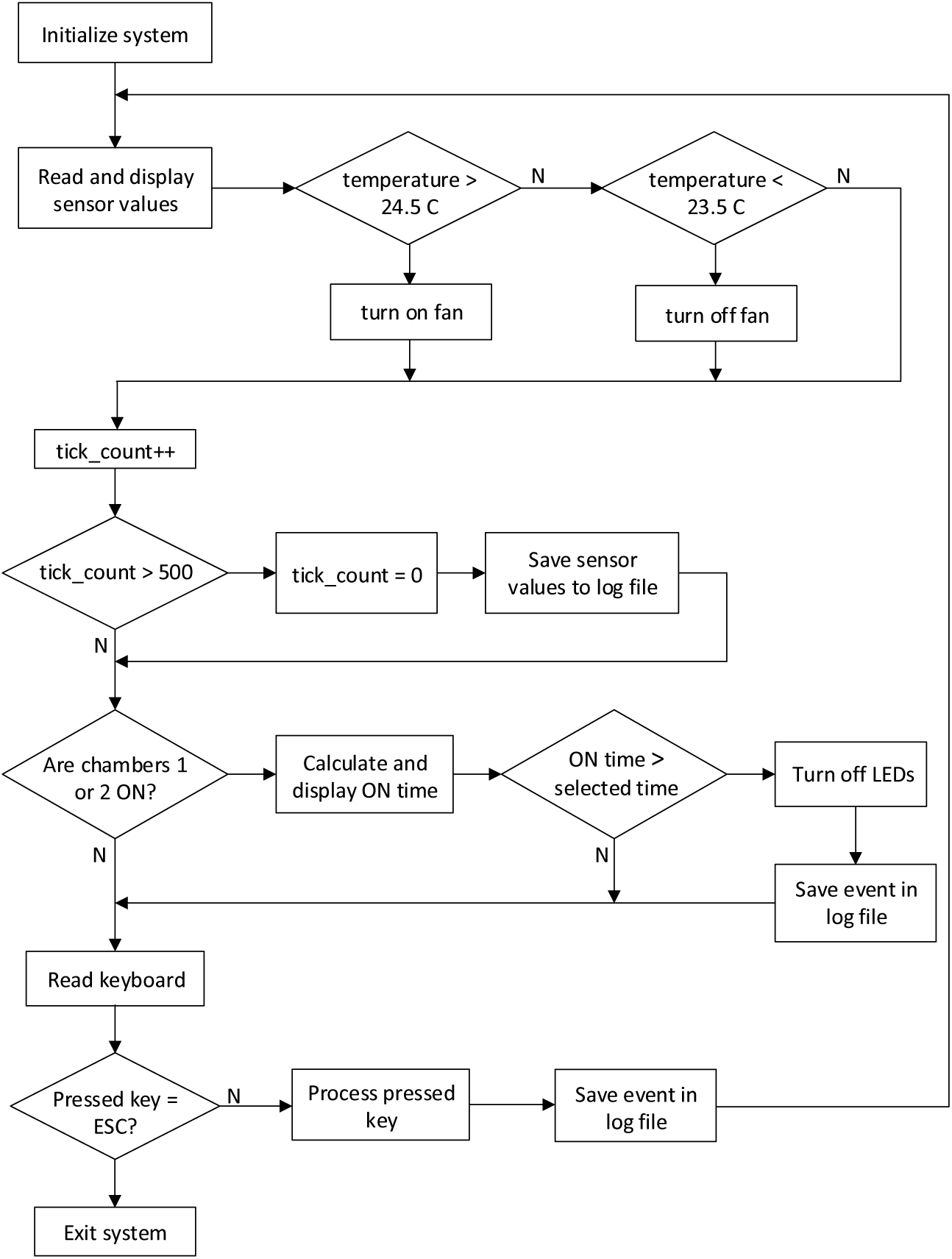
Flow diagram of computer program. The computer program consist of a main loop that reads and displays sensors values and turns off the LEDs if their on time has exceeded the selected stimulation time. The program also implements an ON/OFF temperature control.

## 3. Design files

Table 1 lists the provided files that are required to build and operate the Drosophila Optical Stimulator.

**Table 1:** Design files

| Design file name | File type | Open source license | Location of the file |
| --- | --- | --- | --- |
| LED_board.zip | ZIP file | CERN OHL v1.2 | Google Drive |
| Pi2GPIO_board.zip | ZIP file | CERN OHL v1.2 | Google Drive |
| regulator_board.zip | ZIP file | CERN OHL v1.2 | Google Drive |
| STL.zip | ZIP file | CERN OHL v1.2 | Google Drive |
| DXF.zip | ZIP file | CERN OHL v1.2 | Google Drive |
| code.zip | ZIP file | Three-clause BSD | Google Drive |
| bill_of_materials.osd | OSD file | None | Google Drive |

The following is a brief summary of the content of these files:

1. LED_board.zip: contains the Gerber files needed to fabricate the LED board.
2. Pi2GPIO_board.zip: contains the Gerber files needed to fabricate the Pi2GPIO board.
3. regulator_board.zip: contains the Gerber files needed to fabricate the regulator board.
4. STL.zip: contains the .STL files required to 3D print the parts needed to build the stimulator’s enclosure.
5. DXF.zip: contains the .DXF files required to laser cut the acrylic parts needed to build the stimulator’s enclosure.
6. code.zip: contains the Phyton code that runs on the Raspberry Pi computer and controls the stimulator’s hardware and interfaces with the user.
7. bill_of_materials.osd: is a spreadsheet listing the parts required to build the stimulator.

## 4. Bill of Materials

The bill of materials includes (BOM) over 40 different items. To improve the clarity of this article, the BOM is provided as the accompanying file: `bill_of_materials.ods`. The second column in the BOM is a label that uniquely designates each component. This label is used in the build instructions and schematic diagrams. The BOM also provides a link to online vendors where components can be acquired from. The total cost of parts for the device is USD 757.04.

## 5. Build Instructions

The Drosophila Optical Stimulator is composed of several electronic components housed in a custom-made enclosure. The electronic components are organized in custom and commercially-available electronic boards. There are three custom board designs: LED board, Pi2GPIO board and regulator board. The location of the Gerber files needed to fabricate these boards are provided in Section 3. These boards can be fabricated through a PCB manufacturing company [31]. Sixteen copies of the LED board design are required. One copy of the Pi2GPIO board and the regulator board are needed. After the boards are fabricated, they must be populated with the corresponding electronic components. In each board, a text label and an outline for each electronic component are printed in the silkscreen layer. Use these labels and the outlines to place and solder the electronic components in their right location. The component labels printed on the boards match with text in the **Designator** column in the BOM. For components with polarity such as diodes and electrolytic capacitors, the cathode is highlighted on the component outline.

The commercially-available boards are the Raspberry Pi computer (item 19 in the BOM) and the light and temperature sensors (items 32 and 33 in the BOM). The light and temperature sensors communicate with the embedded computer through an I^2^C serial bus and they need unique addresses to be able to share the common I^2^C bus. Set the address of the light sensor for chamber 1 (LUX1) to 0x49, the address of the light sensor for chamber 2 (LUX2) to 0x39, the address of the temperature sensor for chamber 1 (TEMP1) to 0x19 and the address of the temperature sensor for chamber 2 (TEMP2) to 0x18. Instructions on how to set the addresses of these sensors can be found online [32, 33].

Figure 7 shows a diagram of the electrical connections between the different boards and components of the optical stimulator. In the figure the wires have been color coded to help the reader follow the wires throughout the diagram. Green color wires denote wires carrying data or control signals for chamber 1. Similarly, blue color wires denote wires carrying data or control signals for chamber 2. Black wires connect the ground terminals of the different boards. Red wires carry 12 V, orange wires carry 3.3 V and purple wires carry 5 V. While Fig. 7 shows all the electrical connections as reference. During assembly these connections will have to be made in a particular order as explained below.

**Figure 7:**
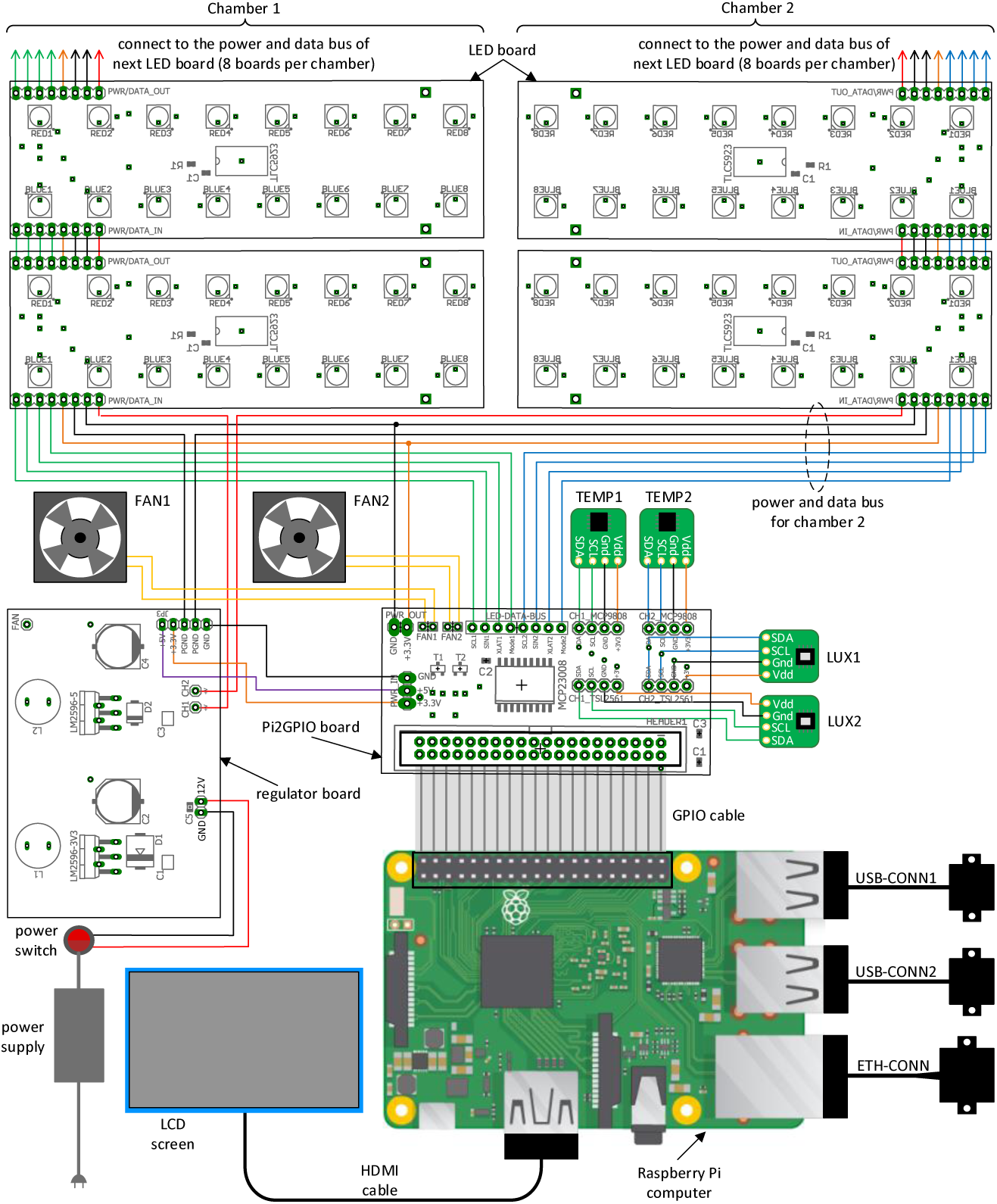
Electrical connection diagram.

Building the Drosophila Optical Stimulator can be accomplished in 12 steps (see Fig. 8 for visual reference). The enclosure for the stimulator is built from 3D printed parts and laser-cut acrylic parts. Each 3D printed part is printed from a .stl file. The acrylic parts are cut out from acrylic panels using a laser cutter according to the shapes specified in .dxf files.

**Figure 8:**
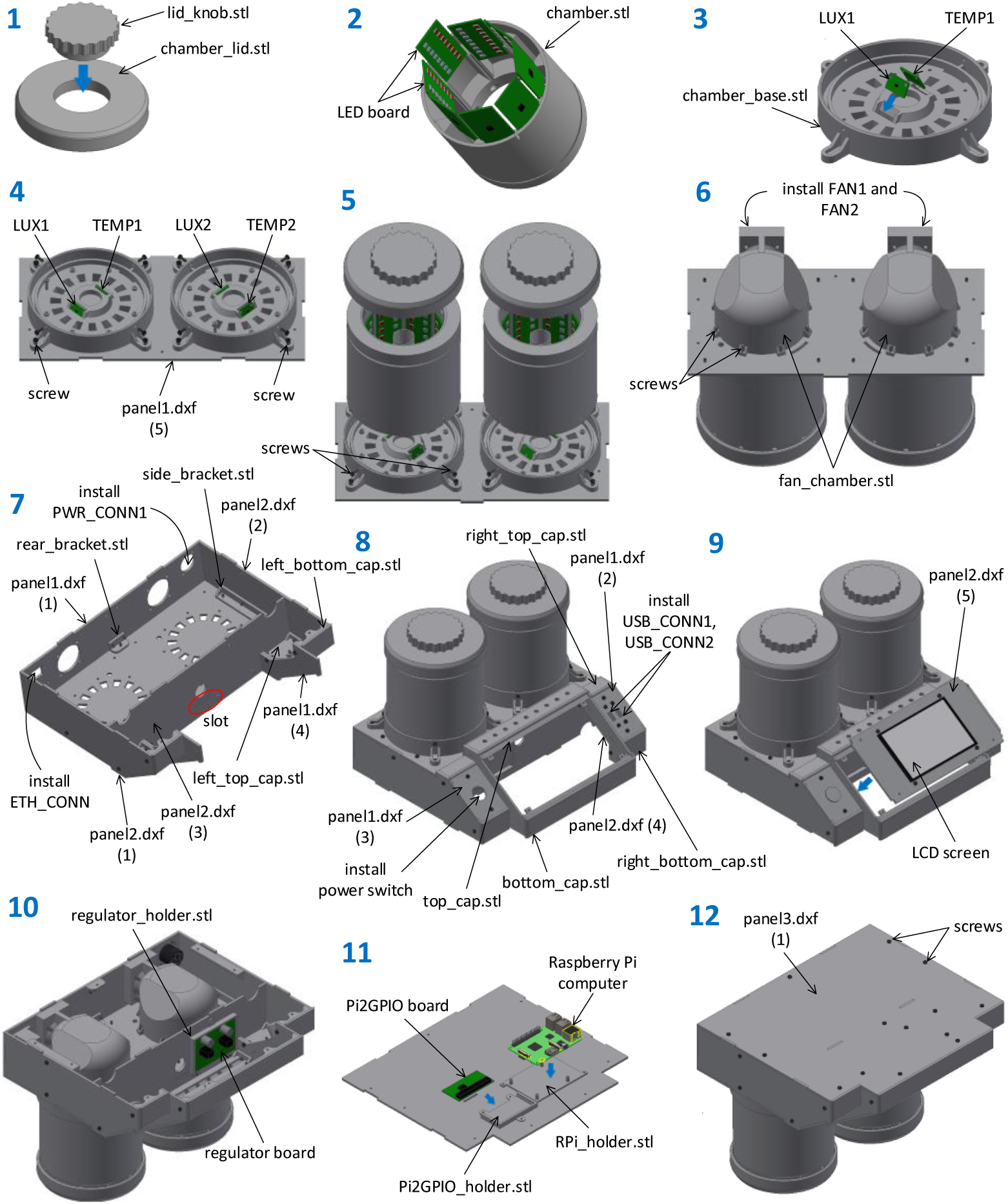
Assembly steps. The enclosure is built from 3D printed parts as well as laser-cut acrylic parts. Each 3D printed part is printed from the indicated .stl file. The acrylic parts come from one of three acrylic panels that are laser cut according to the shapes specified in the .dxf files.

1. Fabricate the chamber lid. A lid is made out of two parts: the knob (`lid_knob.stl`) and the lid body (`chamber_lid.stl`). Print out these two parts and insert the knob into the lid body (see Fig. 8 (1)). Repeat this step to fabricate lids for both chambers.
2. Print the chamber body (`chamber.stl`). Connect the power and data buses of the LED boards in a daisy chain fashion using 24 AWG hook-up wire (items 37 to 39 in the BOM) and a solder iron. Keep the length of the wires between each board to approximately 2.0 cm. The length of the wires on the last board of the chain should be kept at 20 cm for connection to the Pi2GPIO board later on. Inset the wired LED boards into the slots in the chamber body as shown in Fig. 8 (2). Repeat this step to fabricate the bodies for both chambers.
3. Print the chamber base (`chamber_base.stl`). Solder wires to the SDA, SCL, Gnd and Vdd terminals of the temperature and light sensors (LUX1 and TEMP1 for chamber 1 and LUX2 and TEMP2 for chamber 2). The length of these wires should be 20 cm for connection to the Pi2GPIO board later on. Insert the sensor boards in the corresponding slots in the chamber base as shown in Fig. 8 (3). Repeat this step to fabricate the bases for both chambers.
4. Screw the chamber bases to the acrylic support plate using screws and locknuts. The cut-out shape for the acrylic support plate is piece 5 in `panel1.dxf`. Figure 9 (4) shows the cut out shapes for the acrylic panels. The numbers identify a piece within a panel. The .dxf files can be loaded to a laser cutter. Acrylic panels are item 21 in the BOM.
5. Mount the chamber bodies into their corresponding bases and insert screws as shown in Fig. 8 (5).
6. Print the fan chambers (`fan_chamber.stl`) and attach them to the acrylic support plate using screws and locknuts. Solder 20 cm wires to the fans FAN1 and FAN2 (item 29 in the BOM) and install them on the fan chamber using screws.
7. Print a copy of the rear bracket (`rear_bracket.stl`) and two copies of the side bracket (`side_bracket.stl`). Attach the rear wall (piece 1 in `panel1.dxf`) using the rear bracket screws and locknuts as shown in Fig. 8 (7). Install the side wall (piece 1 in `panel2.dxf`) using the side brackets, screws and locknuts. Insert the front wall (piece 3 in `panel2.dxf`) by aligning its slot with the boss in the support plate. Print the parts in files `left_top_cap.stl` and `left_bottom_cap.stl` and install them along with piece 4 from `panel1.dxf` using screws and locknuts as shown in the figure. For clarity, the fan chambers, that should have already been installed, are not shown in the figure. Install the power (PWR CONN1) and the Ethernet (ETH CONN) connectors securing them with screws.
8. Print the parts in files `right_top_cap.stl` and `right_bottom_cap.stl` and install them along with piece 2 from `panel2.dxf` using screws and locknuts. Install plexiglass piece 3 from `panel1.dxf` and plexiglass piece 2 from `panel1.dxf`. Print `top_cap.stl` and `bottom_cap.stl` and install them as shown in Fig. 8 (8). Install the panel-mount USB connectors USB CONN1 and USB CONN2 securing them with screws. Install the power switch as shown in the figure. Wire the power switch to the connector PWR CONN1 installed in step 7.
9. Using screws and locknuts, attach the LCD screen to plexiglass piece 5 in `panel2.dxf` and then screw the plexiglass piece to the rest of the device.
10. Print the regulator board holder from file `regulator_holder.stl` and screw it to the device body as shown in Fig. 8 (10). Solder the wires that go from the regulator board to the Pi2GPIO board and to the LED boards. Mount the regulator board onto its 3D printed holder by press fitting it into the holder.
11. Print the holders for the Pi2GPIO board (`Pi2GPIO_holder.stl`) and the Raspberry Pi computer (`RPi_holder.stl`) and screw them to the device body as shown in Fig. 8 (11). Solder the wires that go from the Pi2GPIO board to the LED boards and the light and temperature sensors. Connect the Pi2GPIO board and the Raspberry Pi computer using the GPIO cable. Insert the SD card with pre-installed Linux OS (item 40 in the BOM) in the SD card slot of the Raspberry Pi computer. Mount the Pi2GPIO board and the Raspberry Pi computer onto their 3D printed holders by press fitting them into the holders. Connect the other ends of ETH_CONN, USB CONN1 and USB_CONN2 to the Ethernet port and USB ports of the Raspberry Pi computer. Connect the power input of the LCD screen to one of the available USB connectors on the Raspberry Pi computer using the USB cable (item 36 in the BOM).
12. Screw the bottom panel (piece 3 from `panel3.dxf`) to close the device. Cut off the output jack from the power supply (item 28 in the BOM) and solder in its place the connector PWR CONN2. Inset PWR CONN2 into PWR CONN1. Connect a USB keyboard and a mouse to the USB connectors on the front of the stimulator.

**Figure 9:**
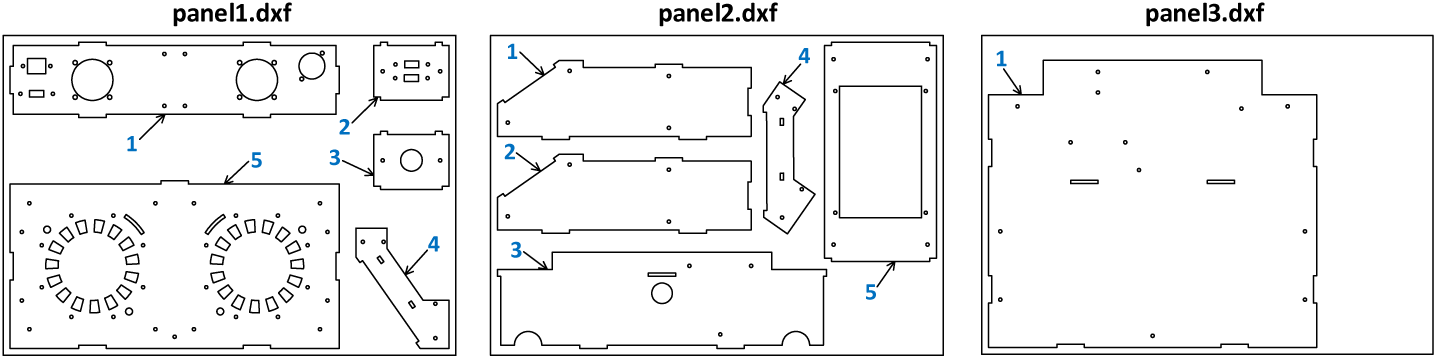
Cut out shapes for the acrylic panels. The numbers identify a piece within a panel. The .dxf files can be loaded to a laser cutter. Acrylic panels are item 21 in the BOM.

## 6. Operation Instructions

Before using the stimulator we need to install support libraries that will allow the application software to access the GPIO pins and the I^2^C of the Raspberry Pi. Connect the device to an Ethernet network with Internet access and power up the stimulator by flipping the power switch. The LCD screen should show messages as the Raspberry Pi computer boots up. Open a Terminal window and install the Adafruit Pi Python repository, the Rpi.GPIO library, the i2c-tools utility and the i2c support for the ARM core and Linux kernel. Refer to [34] for a step-by-step guide on how to install these support libraries and tools. To verify that the I^2^C devices are working properly, issue the following command from the Terminal window:

~~~
sudo i2cdetect -y 1
~~~

A list of the addresses of all the I^2^C devices connected to the Raspberry Pi computer will be displayed. The following addresses should be listed: 49, 19, 39, 18 and 27. If any of these addresses are missing from the list check the wiring that connects the different electronic boards. Unzip the provided `code.zip` file and copy its contents to the `home` directory of user `pi` (default user) on the Raspberry Pi computer. Create a folder in the `home` directory called `logs`. This folder can be created by issuing the following command from a Terminal window:

~~~
mkdir logs
~~~

Log files will be created in this directory. The name of the log files have the following structure: `log_yyyy_mm_dd_hh_mm.txt`, where `yyyy` will be replaced by the year, `mm` by the month, `dd` by the day, `hh` by the hour and `mm` by the minute the stimulator control program was started. To run the stimulator control program, issue the following command from the Terminal window:

~~~
sudo python chamber.py -a
~~~

The screen shown in Fig. 5 will appear. To set stimulation parameters, such as stimulation time, light intensity, color and to turn on or off the LEDs, press 1 for chamber 1 or 2 for chamber 2. A menu will appear on the lower part of the screen. Follow the menu instructions to set the stimulation parameters. The light and sensor readings for both chambers are displayed on the screen at all times. When the LEDs are on, the sensors readings are recorded on the log file every 30 seconds. This interval can be changed by passing its value in seconds as an argument to the control program when it is started. For instance, to record the sensors readings every 5 seconds, start the control program in the following manner:

~~~
sudo python chamber.py -a -t 5
~~~

To operate the stimulator control program remotely, log in to the Raspberry Pi computer using a Secure Shell (SSH) client and issue the commands shown above. Remote login requires knowledge of the IP address of the Raspberry Pi computer. To find out the IP address, type: `sudo ifconfig` and then press Enter on the Terminal window.

## 7. Validation and Characterization

Figure 10 shows the fabricated Drosophila Optical Stimulator. The stimulator was characterized and used to assess blue light-induced retinal degeneration in *Drosophila*. This section presents the characterization and experimental results.

**Figure 10:**
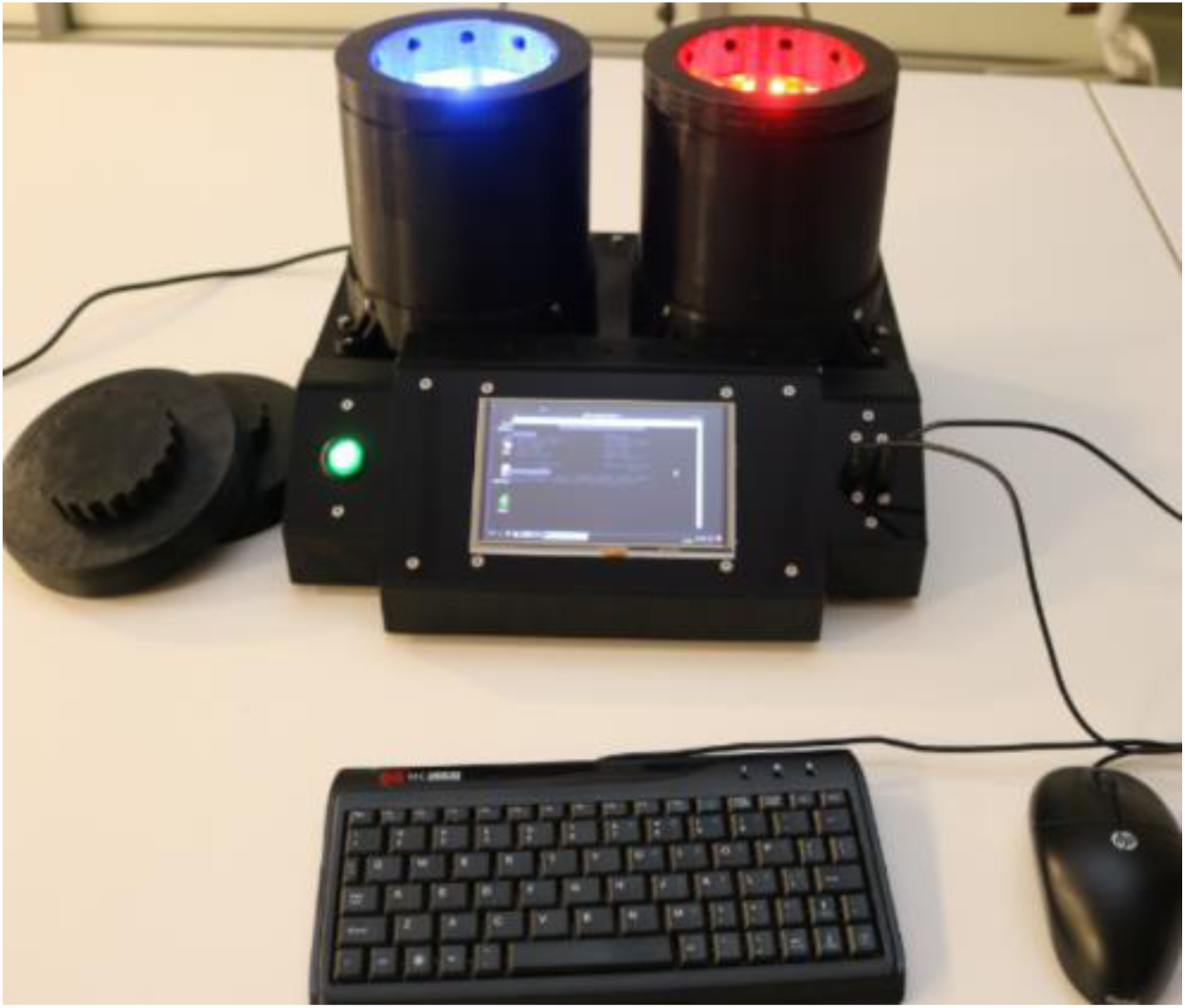
Fabricated optical stimulator. The prototype was fabricated, tested and employed to assess blue-light induced retinal degeneration in *Drosophila* flies.

In the first characterization test the temperature behavior of the stimulator was evaluated for different LED current levels. Figure 11 shows the output of the light and temperature sensors in the left chamber when the blue LEDs are turned on at 60% of maximum LED current *I_max_*. Notably, in the first three minutes the temperature increases linearly as the heat generated by the LEDs and the electronic circuits warm up the air inside the chamber. When the temperature reaches 24.5 °C the fan is turned on and the temperature increase slows down until it settles at around 26 °C. The LED illuminance decreases at the beginning as the LEDs warm up which is an expected behavior in LEDs. However, the illuminance stabilizes as the temperature in the chamber reaches a steady-state value.

**Figure 11:**
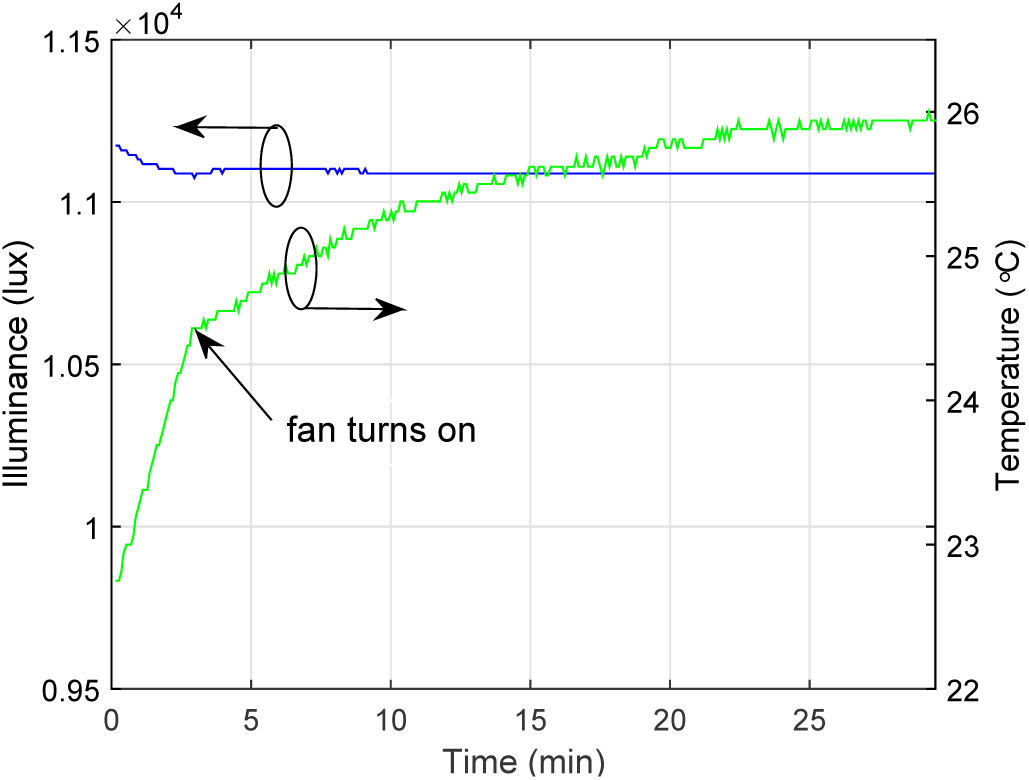
Acquired illuminance and temperature waveforms. The waveforms were acquired from the sensors in the left chamber for a 60% light intensity of the blue LEDs.

The steady-state temperature value was recorded for each chamber as the LED current was varied from 20% to 100% of *I_max_*. The measurements were repeated three times for each LED current level to verify the repeatability of the measurements. Before each repetition the chambers were allowed to cool down to room temperature. Figure 12 shows the average steady-state temperature over the three repetitions as the height of the bars for both the left and right chambers and for blue and red light. The error bars denote the maximum and minimum steady-state temperature over the three repetitions. The average temperature reaches a maximum of 28.17 °C at 100% LED current. Although this temperature is a bit higher than the value established in the design requirements, is not of sufficient magnitude to induce heat shock in the flies.

**Figure 12:**
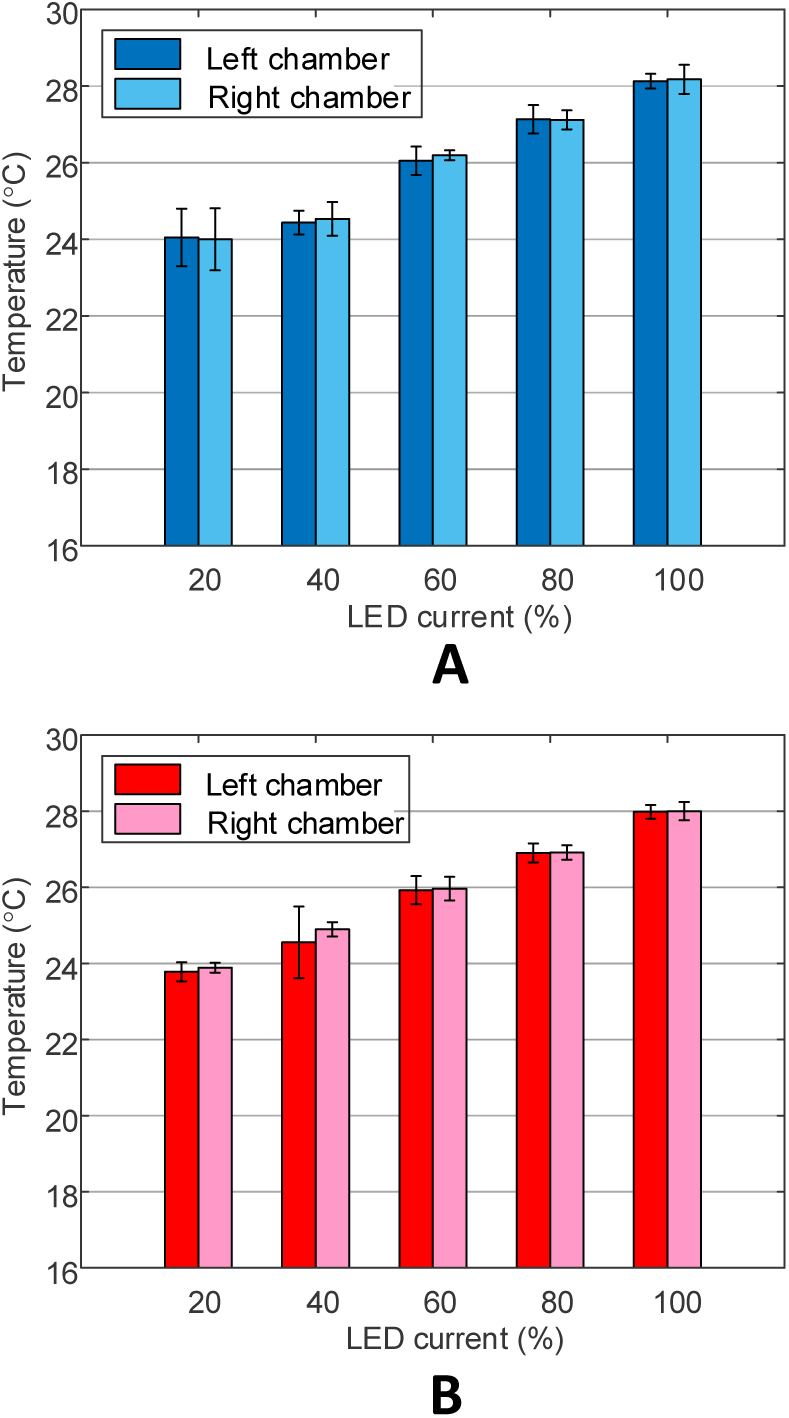
Measured steady-state chamber temperature for different LED current percentage values. The height of the bars denote the average steady-state chamber temper ature over three repetitions. The errors bars represent the maximum and minimum steady-state temperature values over the three repetitions. (A) Blue LEDs. (B) Red LEDs.

Figure 13 shows the illuminance inside the chambers for blue and red as measured by the light sensors as the LED current is varied from 0% to 100% of *I_max_*. Figure 13 (A) shows the illuminance for the blue LEDs while Fig. 13 (B) shows the illuminance for the red LEDs. Notably, there is an almost linear relationship between the illuminance and the current through the LEDs. The illuminance read by the left chamber sensor is consistently lower than the illuminance read by the right chamber sensor when the red LEDs are on (Fig. 13 (B)). This difference is due to different sensitivity to red light between the photodiodes in the light sensors. To verify this difference the optical power inside the chambers was measured as a function of LED current using the S120C calibrated optical power sensor from Thorlabs. The measured optical power was divided by the power sensor detector area to obtain the power density. Figure 13 (C) and (D) show the optical power density inside the chambers for blue light (C) and for red light (D). Notably, there is no significant difference between the power density levels of the red LEDs in the left and right chambers. Using this information the light sensors inside the chambers could be calibrated to provide consistent illuminance readings for red light.

**Figure 13:**
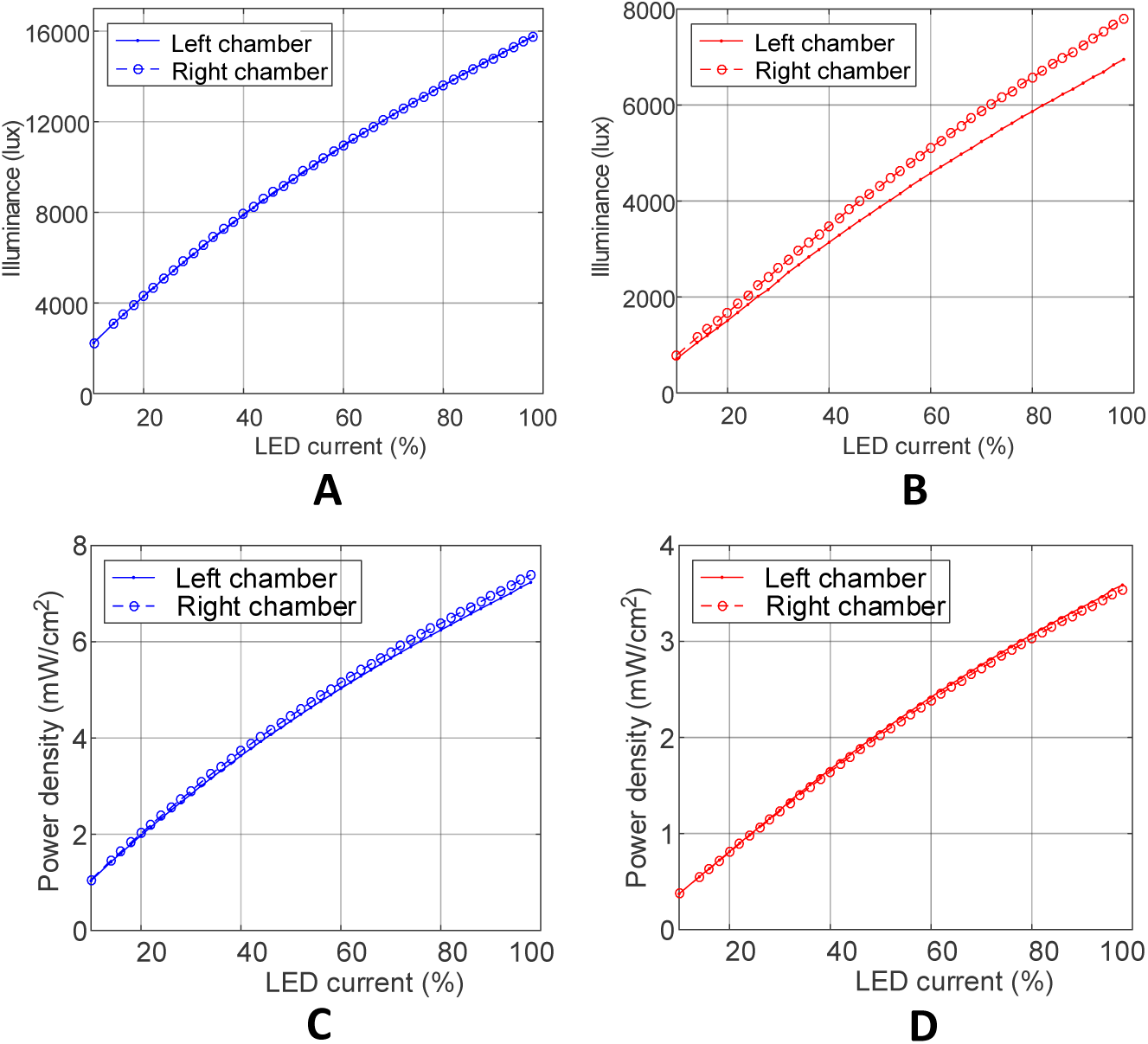
Measured illuminance and optical power density for different LED current percentage values in both chambers. Illuminance was measured with the light sensors embedded in the stimulator. Optical power was measured with a calibrated power meter. The optical power was divided by the area of the power meter detector to obtain power density. (A) Illuminance vs LED current for the blue LEDs. (B) Illuminance vs LED current for the red LEDs. (C) Power density vs LED current for the blue LEDs. (D) Power density vs LED current for the red LEDs.

The stability of the device during long-duration experiments was also evaluated. To this end, three 8-hour stimulation sessions were performed on three different days with the same stimulation parameters: blue light at 42% LED current. These stimulation parameters are typical for the biological experiments that have been performed with the device. The maximum illuminance change was 0.18% over the three different stimulation sessions. Hence, the fabricated optical stimulator is able to provide stable light stimulation levels over long periods of time.

### 7.1. Biological Experiments

The fabricated programmable optical stimulator was used to establish a protocol for assessing blue light-induced retinal degeneration in young flies (*Drosophila melanogaster*). To do this, we used male *w*^1118^ flies that lack eye pigment but are otherwise wild type, and are a standard model system for studying visual function and retinal degeneration. We collected newly eclosed (hatched from pupal cases) male flies and aged these for 6 days in 12 hours of light (~15 lux, fluorescent)/12 hours dark at 23 – 25°C in polystyrene 25 × 99 mm vials (VWR, #89092-722) on standard cornmeal-yeast fly food (Fig. 14(A)). Flies were then separated into three experimental groups that each consisted of 5 male flies. The first group was stimulated with blue light for 8 hours at an illuminance of 7949 lux. The second group was stimulated with red light for 8 hours at an illuminance of 7994 lux. The third group was kept in the dark for 8 hours as a negative control for light exposure. Following each light or dark treatment, flies were incubated in the dark for 12 days to enable sufficient time for rhabdomeres to degenerate so that photoreceptor loss (retinal degeneration) could be accurately assessed. Photoreceptor loss was analyzed using confocal microscopy of retinas immunostained with antibodies against Rhodopsin 1 and with phalloidin, which stains F-actin in the rhabdomeres of each photoreceptor cell. To assess retinal degeneration, we count the number of intact R1 – R6 rhabdomeres as identified by phalloidin and Rhodopsin 1 staining in the retina of flies from each experimental group. In this experiment, the loss of a rhabdomere indicates that corresponding photoreceptor cell has died and that retinal degeneration is occurring.

**Figure 14:**
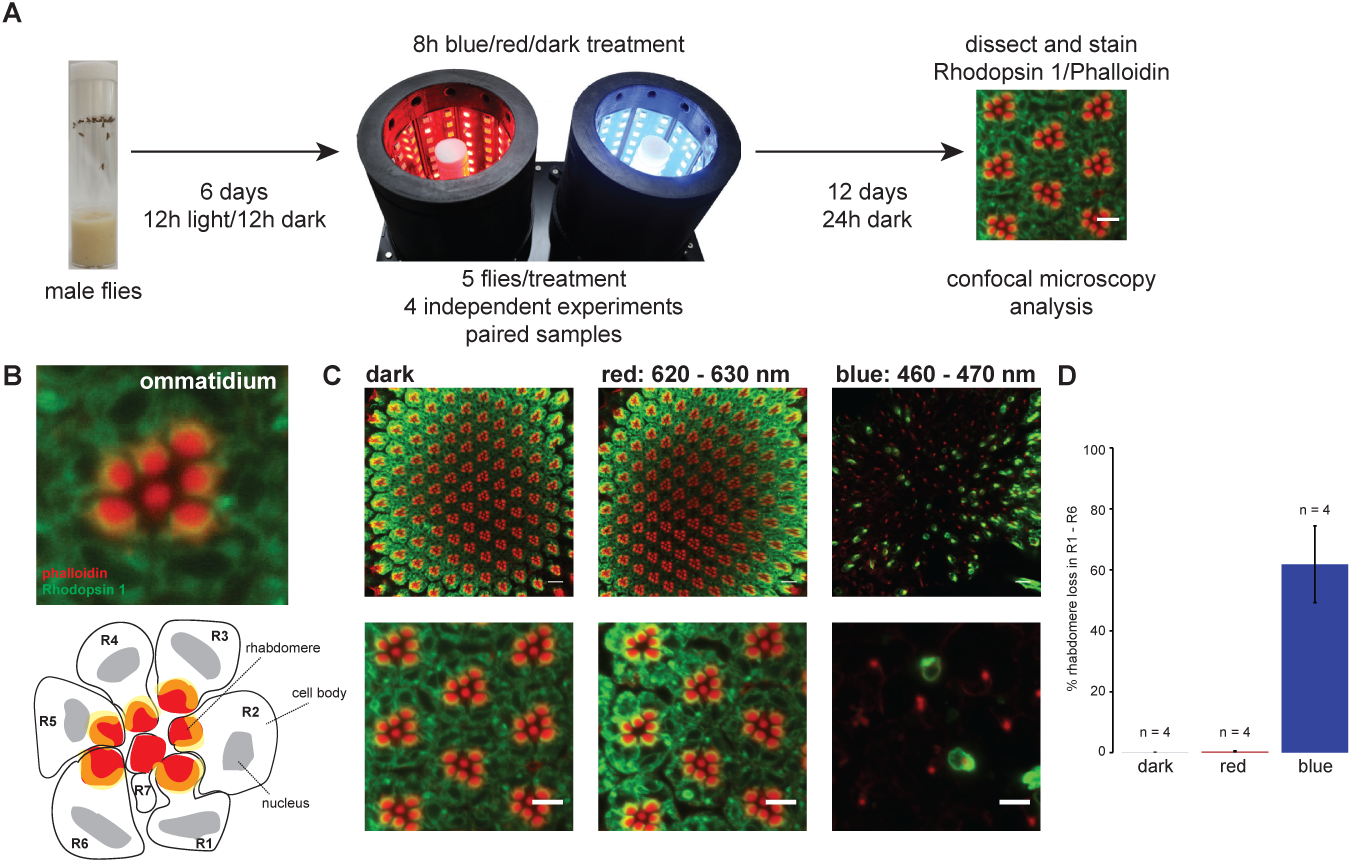
Biological experimental results. (A) Schematic of the experimental design. Newly eclosed mated *w*^1118^ male *Drosophila melanogaster* were raised in 12 h/12 h light/dark conditions for 6 days prior to treatment with either 8 h or red or blue light at ~8000 lux relative to dark control. Flies were incubated in the dark for 12 days post-treatment, following which retinas were immunostained with antibodies against Rhodopsin 1 (Rh1, green) and F-actin (phalloidin, red) and visualized using confocal microscopy. (B) A confocal microscopy image of a single ommatidium stained as described in panel A is presented in the upper panel, and a schematic representation of the photoreceptor neurons (R cells) present in this image is shown in the lower panel for comparison. (C) Representative images from single flies exposed to 8 h of red or blue light at ~8000 lux relative to the dark control. Confocal microscopy images (single plane) of retinas stained with Rh1 and phalloidin at low magnification (upper panel) and high magnification (lower panel) are shown. (D) Bar plot showing quantification of the retinal degeneration phenotypes illustrated in panel C. Photoreceptor loss for R1 – R6 cells was determined by analyzing rhabdomere loss in retinas from 5 individual eyes from 5 independent animals in 4 independent matched biological experiments (n, biological experiment). Data is shown as mean ± standard deviation.

The method for immunostaining retinas is described briefly as follows. Retinas were dissected and fixed in phosphate-buffered saline (1XPBS, pH 7.4) containing 4% paraformaldehyde, 1 mM EGTA, 1 mM MgCl_2_ for 25 min, washed 4 times in PBST (PBS containing 0.03 % (v/v) Triton X-100), and incubated with anti-rh1 (4C5, mouse, 1:50, Developmental Studies Hybridoma Bank) and phalloidin (A22287, 1:100, Thermo Fisher Scientific Inc.) for 8 – 16 hours at 4°C. Eyes were washed 4 times in PBST, incubated with fluorophore-conjugated secondary antibody (A21202, Thermo Fisher Scientific Inc.) for 8 – 16 hours at 4°C, washed 4 times in PBST, and mounted in 2XPBS containing 70% sorbitol (w/v) (S1876, Sigma-Aldrich Co.). Laser scanning confocal imaging was performed using a Zeiss LSM710 confocal microscope. Images were collected using Plan-Apochromat 63×/1.4 Oil at Z-stack step size of 1 *μ*m. Rhabdomere loss was quantified using the stacked images to assess retinal degeneration.

Figure 14(A) shows the experimental protocol used to assess blue light-induced retinal degeneration in flies as outlined above. Figure 14 (B) shows a single ommatidium stained for Rhodopsin 1 and phalloidin, which marks F-actin. Whereas phalloidin labels the rhabdomere in all photoreceptors R1 – R8, Rhodopsin 1 (shown in green) only labels a subset of these photoreceptors, R1 – R6. A characteristic crescent of Rhodopsin 1 protein can be observed at the base of each rhabdomere in these photoreceptor cells (yellow due to the overlap of F-actin and Rhodopsin 1 signals). The central R7 and R8 photoreceptor neurons do not express Rhodopsin 1, and R8 cells are present in a deeper layer of each ommatidium so are not visible in the single plane images shown in this figure. The dark regions inside each R1 – R6 neuron correspond to the nucleus of each cell. The position of each photoreceptor cell, rhabdomere and nucleus are indicated below the confocal image of the immunostained ommatidium to orient the reader.

To test if the blue light produced using the fabricated optical stimulator induces retinal degeneration in flies treated using our protocol, we examined rhabdomere loss as an indicator of photoreceptor loss, and therefore retinal degeneration. When flies were exposed to 8 hours of blue light at 7949 lux, equivalent to 20.12 log quanta/cm^2^, we consistently observed high levels of retinal degeneration as shown in Fig. 14 (C, blue). Whereas the dark control flies have well organized retinas with intact rhabdomeres in all ommatidia, as demonstrated by phalloidin and Rhodopsin 1 staining, the blue light-exposed flies show strong loss of rhabdomeres across multiple ommatidia in the retina. We quantified this rhabdomere loss by analyzing retinas from 4 independent biological experiments each consisting of 5 individual flies, and showed that 62% of R1 – R6 cells undergo cell death following blue light exposure relative to less than 1% of cells in the dark control (Fig. 14 (D)). To test if the retinal degeneration induced by blue light was specific to this wavelength, or was a consequence of the high intensity used in this experiment, we exposed a second group of flies to red light for 8 hours at an intensity of 7994 lux. Strikingly, the red light-exposed flies do not show any significant loss of rhabdomeres (Fig. 14 (C) and (D)), and have less than 1% of rhabdomere loss, similar to the level observed in the dark control. Based on these data, we conclude that the fabricated programmable optical stimulator induces retinal degeneration in flies treated with blue light using our protocol, and that this retinal degeneration results from the wavelength rather than intensity of light used.

## 8. Conclusions

A programmable optical stimulator for *Drosophila* fly eyes has been presented. The stimulator uses LEDs and an embedded computer to control illuminance, wavelength (blue or red) and duration in two independent chambers. Furthermore, the stimulator is equipped with per-chamber light and temperature sensors and a fan to monitor light intensity and to control temperature. An ON/OFF temperature control implemented on the embedded computer keeps the temperature inside the chambers around 28°C to avoid heat shocking the flies. A custom enclosure was fabricated to house the electronic components of the stimulator, provide a light-impermeable that allows air flow environment and to allow users to easily load and unload fly vials. The performance of the stimulator was characterized. Characterization results show that the fabricated stimulator can produce light at illuminances ranging from 0 to 16000 lux and power density levels from 0 to 7.2 mW/cm^2^ for blue light. For red light the maximum illuminance is 8000 lux which corresponds to a power density of 2.54 mW/cm^2^. The fans and the ON/OFF temperature control are able to keep the temperature inside the chambers below 28.17°C. Finally, biological experiments with white-eye flies were performed to assess the ability of the fabricated device to induce blue light retinal degeneration. Retinal degeneration was observed in flies exposed to 8 hours of blue light at 7949 lux. However, flies exposed to red light for the similar duration and light intensity do not show retinal degeneration.

## Author’s contributions

XC performed the characterization tests and biological experiments, WDLS designed the electronic hardware, wrote the software and wrote the paper, TZ designed the enclosure and built the programmable stimulator, DR conceived the original single-LED prototype and provided consultation during the experiments, VMW designed and oversaw the biological experiments.

## Acknowledgements

This work was supported by the Ralph W. and Grace M. Showalter Research Trust. Support from the National Institutes of Health (NIH) R01EY024905 to V. Weake is gratefully acknowledged. NIH Shared Instrumentation Grant, NCRR 1 S10 RR023734-01A1 supported the Zeiss LSM 710 confocal microscope. X. Chen was supported in part by NIH grant R01EY10306.

